# Mucosal-associated invariant T (MAIT) cells mediate protective host responses in sepsis

**DOI:** 10.1101/2020.02.05.935866

**Authors:** Shubhanshi Trivedi, Daniel Labuz, Cole P. Anderson, Claudia V. Araujo, Antoinette Blair, Elizabeth A. Middleton, Alex Tran, Matthew A. Mulvey, Robert A. Campbell, J. Scott Hale, Matthew T. Rondina, Daniel T. Leung

**Affiliations:** Division of Infectious Disease, University of Utah, Salt Lake City, Utah; Division of Pulmonary and Critical Care, University of Utah, Salt Lake City, Utah; Division of General Internal Medicine, Department of Internal Medicine, University of Utah, Salt Lake City, Utah; Division of Microbiology and Immunology, Department of Pathology, University of Utah, Salt Lake City, Utah; George E. Wahlen VAMC Department of Internal Medicine and GRECC, University of Utah, Salt Lake City, Utah; Molecular Medicine Program, University of Utah, Salt Lake City, Utah

## Abstract

Sepsis is a systemic inflammatory response to infection and a leading cause of death. Mucosal-associated invariant T (MAIT) cells are innate-like T cells enriched in mucosal tissues that recognize bacterial ligands. We investigated MAIT cells during clinical and experimental sepsis, and their contribution to host responses. In experimental sepsis, MAIT-deficient mice had significantly increased mortality and bacterial load, and reduced tissue-specific cytokine responses. MAIT cells of WT mice expressed lower levels of IFN-γ and IL-17a during sepsis compared to sham surgery, changes not seen in non-MAIT T cells. MAIT cells of patients presenting with sepsis were significantly reduced in frequency, more activated, and had decreased IFN-γ production when stimulated, compared to healthy donors and paired 90-day post-sepsis samples. Our data suggest that MAIT cells are highly activated and become dysfunctional during clinical sepsis, and contribute to tissue-specific cytokine responses that are protective against mortality during experimental sepsis.

## Introduction

Sepsis is a life threatening syndrome caused by dysregulated host responses to infection resulting in organ dysfunction (1). It is a leading cause of death worldwide, accounting for 5.3 million deaths annually (2). Sepsis is characterized by a systemic acute pathologic hyper-inflammatory response and an opposing anti-inflammatory response that can occur concurrently (3). Hallmarks of sepsis include a decreased ability to eliminate primary pathogens and an increased susceptibility to secondary nosocomial infections (4). In addition, it is associated with a protracted immunosuppressed state that contributes to morbidity and mortality (5). Unfortunately, clinical trials over the past 3 decades of interventions targeting the hyper-inflammatory state of sepsis have largely failed to consistently improve clinical outcomes (6-8), and effective treatments are needed.

Many components of host responses are altered during sepsis, including activation of innate effector cells such as platelets, neutrophils, epithelial cells, and endothelial cells. In some settings, cellular activation is associated with dysfunctional responses and adverse clinical outcomes. Studies on immunosuppression in sepsis have largely focused on exhaustion, apoptosis, and reprogramming of a broader range of effector cells, including T cells, B cells, neutrophils, and other antigen-presenting cells (9). Unconventional T cells, such as invariant NKT (iNKT) cells, γδ T cells, and mucosal-associated invariant T (MAIT) cells, bridge the innate and adaptive arms of the immune response, and have the capacity to contribute to both early-phase inflammation and late-phase immunosuppression.

MAIT cells are innate-like T cells restricted by MHC-related molecule 1 (MR1) and MAIT ligands belong to a class of transitory intermediates of the riboflavin synthesis pathway (10). They express an invariant T cell receptor (TCR) TRAV1-2 (or Vα7.2) in humans and TRAV-1 (or Vα19) in mice and a variable, but restricted, number of TRAJ and TCR β chains. MAIT cells are abundant in mucosal tissues such as the liver, lung, and mesenteric lymph nodes and gut lamina propria. In humans, they constitute 1-10% of peripheral blood T lymphocytes, up to 10% of intestinal T cells, and up to 40% of T cells in the liver (11, 12). Recent studies have highlighted the protective role of MAIT cells in host antibacterial responses *in vivo* (13-15). It has also been shown that in patients with sepsis, MAIT cell frequencies are decreased in circulation compared to healthy control donors and uninfected critically-ill patients (16). Septic patients with persistent MAIT cell depletion also have a higher incidence of secondary, ICU-acquired infections (16). However, detailed phenotypic and functional MAIT cell changes during sepsis, as well as the mechanisms by which MAIT cells contribute to host immune responses in sepsis, are not known. In this work, we used complementary, longitudinal studies in sepsis patients and in a relevant murine sepsis model to study the role of MAIT cells in sepsis pathology. We examined immune responses in C57BL/6 wild type (WT) or MR1 knock-out (MR1^-/-^; i.e. MAIT depleted) mice using the cecal ligation and puncture (CLP) model of polymicrobial sepsis. Additionally, we evaluated the phenotype and function of human MAIT cells during acute sepsis and at 3 months after sepsis.

## Results

### MAIT cells have decreased expression of effectors during experimental sepsis

To assess MAIT cell effector function during sepsis, we examined MAIT cells in the lungs of WT mice undergoing CLP, a polymicrobial model of sepsis, or sham surgery. We chose lungs given the frequent association of lung injury with clinical sepsis (17, 18), and because it is among the most abundant site of MAIT cells in mice (19). While we noted similar frequencies of MR1-5OP-RU-tetramer^+^ (from here on, referred to as MR1-tetramer^+^) MAIT cells in CLP mice as in sham mice (Figure 1A and B), we found in sorted MAIT cells that gene expression of MAIT effectors *Ifng* (P = 0.04) and *Il17a* (P = 0.02) were significantly lower (and *GzmB* trended lower, P = 0.14) in septic mice compared to sham mice (Figure 1C). Conversely, in non-MAIT TCRβ^+^CD3^+^ T cell populations, *Ifng* (P = 0.06), *Il17a* (P = 0.16) and *Gzmb* (P = 0.49) mRNA expression levels were similar between septic mice and sham mice (Figure 1D). These data suggest that MAIT cells are specifically impaired during experimental sepsis.

**Figure 1.**
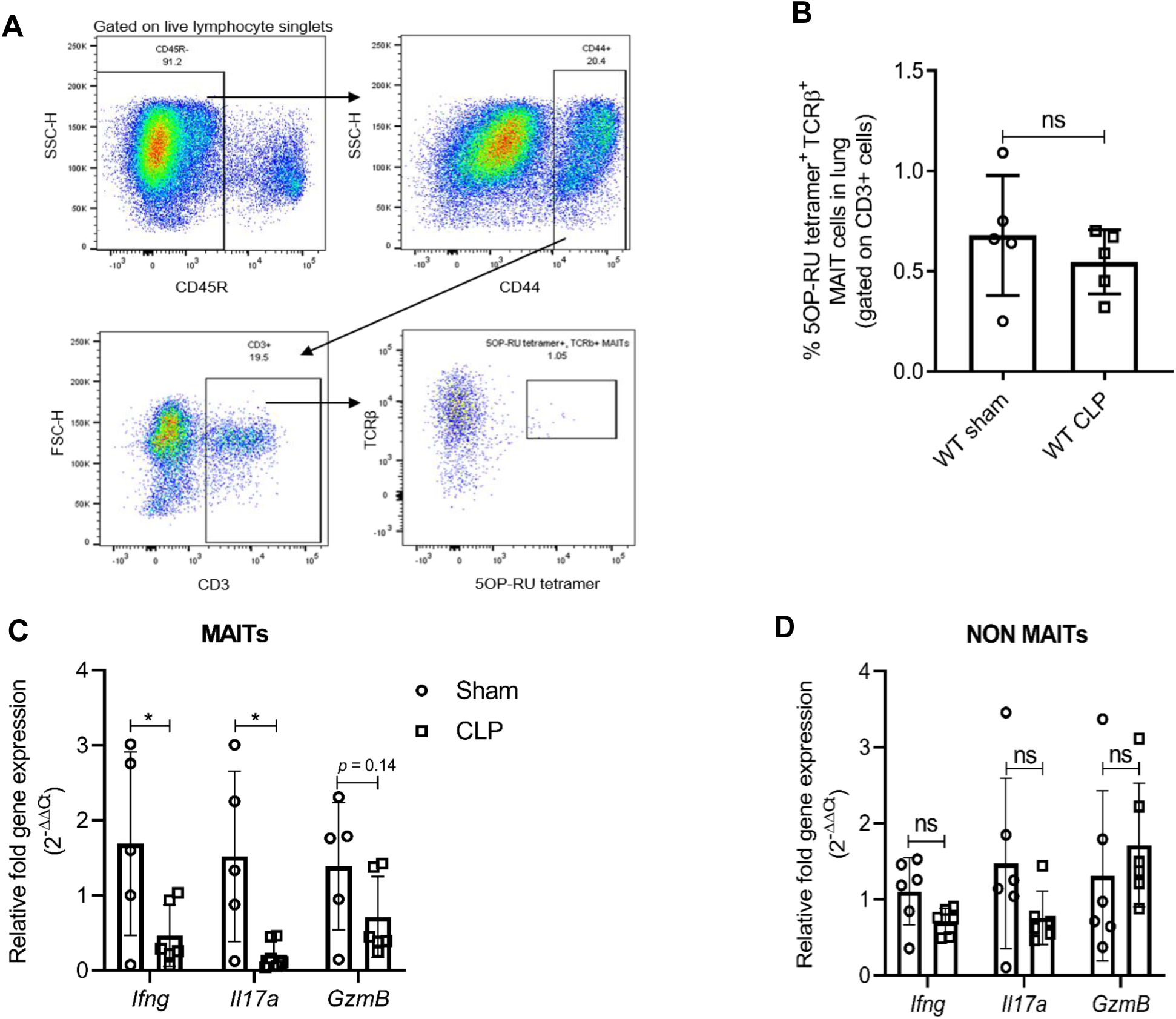
Sepsis induces MAIT-specific changes in inflammatory cytokine expression. (**A**) Gating strategy for isolation of MAIT cells and (**B**) Percentage of TCR^+^ MR1-tetramer^+^ MAIT cells, of CD3^+^ T cells. (**C**) Gene expression in MAIT cells isolated using flow cytometric sorting of homogenized lung tissue after CLP or sham operation (n = 5 per group), as determined by qRT-PCR for *Ifng, Il17a* and *GzmB* genes, (**D**) *Ifng, Il17a*, and *GzmB* mRNA expression in non-MAIT (MR1-tetramer^-^ TCR^+^) CD3^+^ T cell populations. Data are expressed as mean ± SD, and unpaired *t-*test was used for comparisons (data passed Shapiro-Wilk normality test).

### MAIT deficiency increases bacterial burden and mortality during experimental sepsis

We examined whether the absence of MAIT cells altered survival outcomes *in vivo* during experimental sepsis. In the CLP model of polymicrobial sepsis (20, 21), we saw that MR1-deficient mice (MR1^-/-^) which lack MAIT cells, had significantly increased sepsis-related mortality compared to WT mice (Figure 2A). The majority (11/15, ∼73%) of MR1^-/-^ mice died 24 to 48 hours following induction of sepsis, while the majority (13/15, ∼87%) of WT mice survived up to 100 hours after sepsis (Figure 2A). On examining bacterial counts 18 hours following CLP, we found that MR1^-/-^ mice had a significantly higher burden of bacteria in blood compared to WT mice (Figure 2B). To extend these findings to a setting of sepsis due to a single pathogen, we induced sepsis via intraperitoneal injection of a clinical strain of extraintestinal pathogenic *Escherichia coli* (ExPEC). As we observed in the CLP, polymicrobial model of sepsis, MR1^-/-^ mice had significantly higher mortality from ExPEC sepsis, compared to WT mice (Supplementary Figure 1).

**Figure 2.**
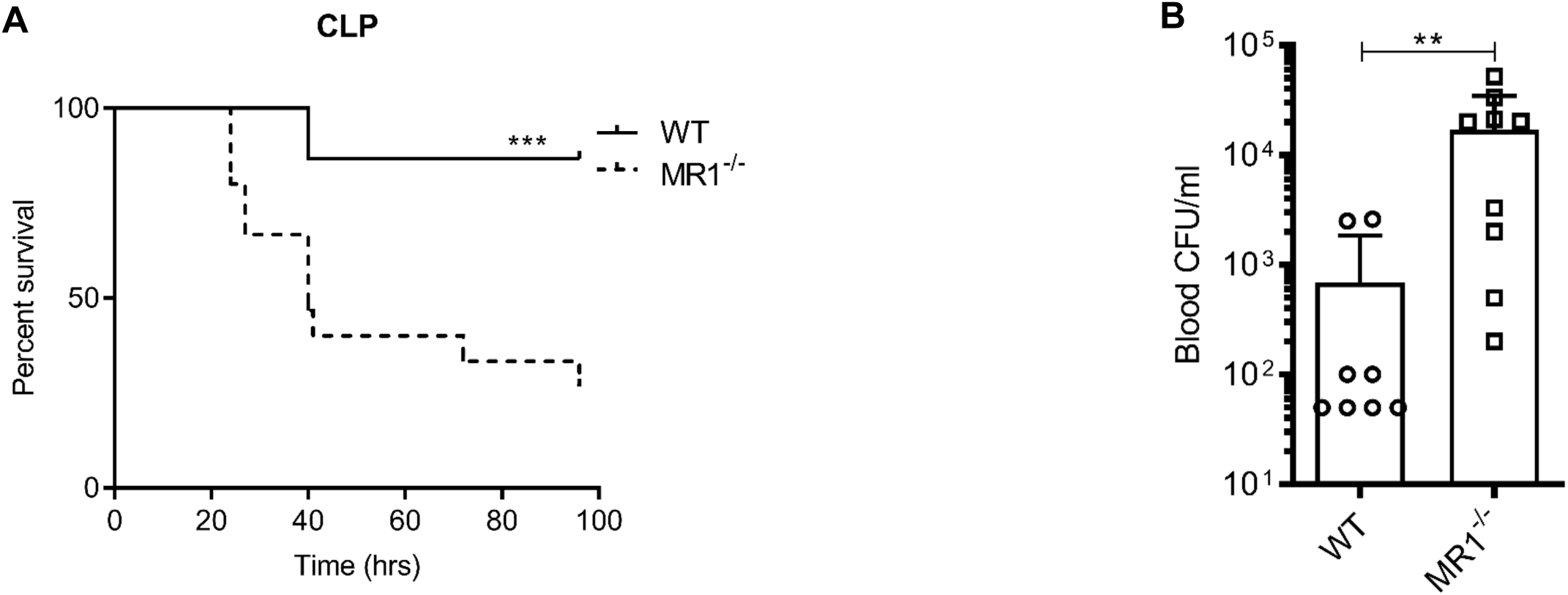
MR1^-/-^ mice had higher sepsis-induced mortality and bacterial burden compared to wildtype mice. (**A**) Age matched MR1^-/-^ and WT mice (n = 15 per group) underwent CLP to induce polymicrobial sepsis. Survival was recorded over a period of 4-5 days (WT versus MR1^-/-^ mice, *** p = 0.0007). Data represents two independent experiments. (**B**) Blood was collected from WT and MR1^-/-^ mice (n = 8-10 per group) 18 h post CLP, and serial dilutions were plated on Luria Broth (LB) agar plates. Colony-forming units (CFU) were determined 24 h after plating and were expressed as CFU/ml. Statistical analysis was performed using Mann-Whitney test.

### MAIT deficiency is associated with reduced tissue cytokine responses in experimental sepsis

To determine the contribution of MAIT cells to sepsis-induced tissue inflammatory responses, we evaluated MAIT-associated cytokines (22) in lung homogenates after CLP. Compared to WT mice, MR1^-/-^ mice had significantly reduced levels of lung IFN-γ, TNFα, IL-17A, IL-10 and GM-CSF: cytokines produced by MAIT cells (22) (Figure 3). Interestingly, serum cytokines were less affected than tissue cytokines in MR1^-/-^ mice following sepsis (Supplemental Figure 2).

**Figure 3.**
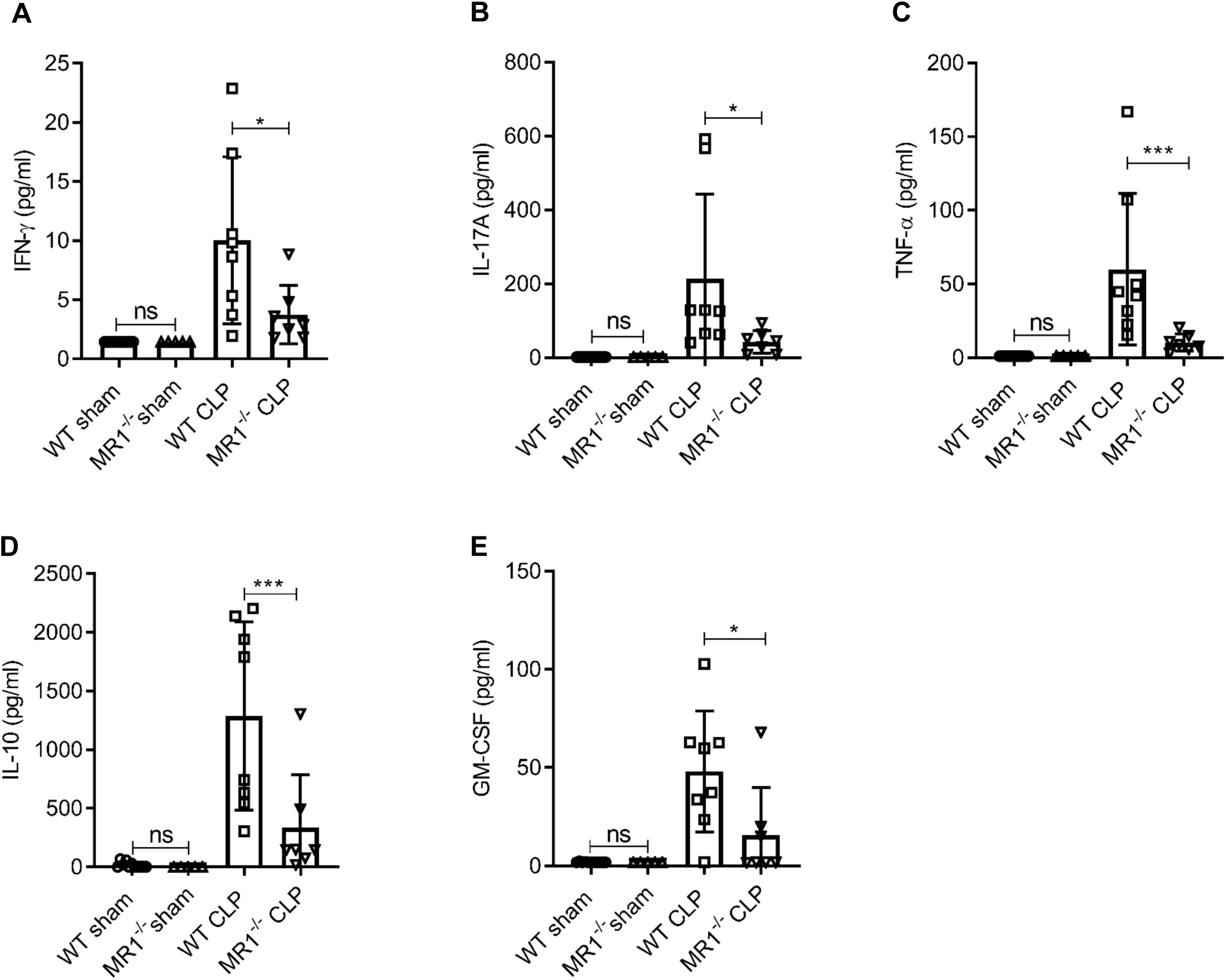
MR1^-/-^ mice had significantly lower levels of cytokines in lung tissue at 18 h post CLP. Levels of (**A**) IFN-γ, (**B**) IL-17A, (**C**) TNF-γ, (**D**) IL-10, and (**E**) GM-CSF were assessed in lungs isolated from WT and MR1^-/-^ mice at 18 h post CLP or sham operation, using a bead-based immunoassay kit. Bars represent mean cytokine levels ± SD. The graphs represent two independent experiment (WT n= 8, MR1^-/-^ n = 7). The statistical significance was determined by the nonparametric Mann-Whitney test.

### MAIT deficiency is associated with reduced tissue-specific interstitial macrophages and monocytic dendritic cell frequencies

MAIT cells can contribute to cytokine release from macrophages (23) and have shown to promote inflammatory monocyte differentiation into dendritic cells during pulmonary infection (24). Thus, we next sought to determine the cause of reduced tissue cytokine production of MR1^-/-^ mice during sepsis. We approached this by determining how an absence of MAIT cells influences the frequencies of lung innate immune cells, including alveolar macrophages (AMs), interstitial macrophages (IMs), monocytic dendritic cells (moDCs) and plasmocytoid DCs (pDCs). As compared to WT mice, MR1^-/-^ mice had a significantly lower proportion of IMs (Siglec-F^-^ CD24^-^ MHCII^+^ CD11c^+^ CD11b^+^ CD64^+^ cells; Figure 4A) and moDCs (Siglec-F^-^ MHCII^+^ CD11c^+^ CD11b^+^ Ly-6C^+^ cells; Figure 4B) following CLP. Loss of MAIT cells did not influence AMs or pDCs, however, following CLP (Figure 4C-D).

**Figure 4.**
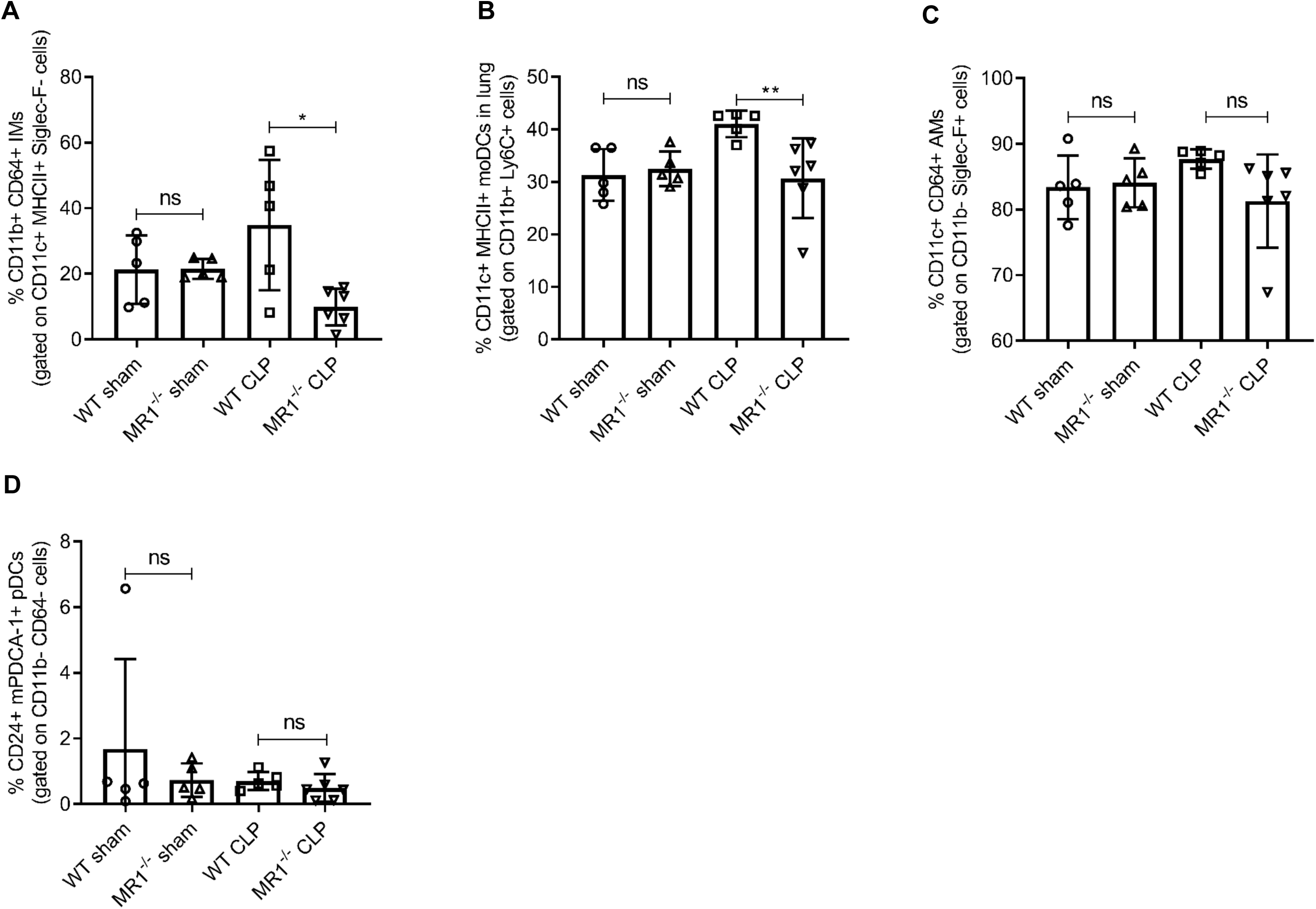
Lower frequencies of interstitial macrophages and monocytic dendritic cells in lungs of MR1^-/-^ mice at 18 h post CLP. Lung cells were obtained from WT (n= 5) and MR1^-/-^ mice (n = 5-6) at 18 h post CLP or sham operation, stained for surface markers and analyzed using flow cytometry. Percentage frequencies of (**A**) interstitial macrophages, (**B**) monocytic dendritic cells, (**C**) alveolar macrophages and (**D**) plasmacytoid dendritic cells were compared between the groups nonparametric Mann-Whitney test. Data were expressed as mean ± SD of two independent experiments.

We next examined whether loss of MAIT cells altered the frequencies of other unconventional T cells such as invariant natural killer T cells (*i*NKT) (25, 26) and γδ T cells (27, 28) in sepsis. We observed that frequencies of *i*NKT and γδ T cells in lungs were reduced during sepsis, consistent with prior studies showing that frequencies of *i*NKT (25, 26) and γδ T cells (27, 28) are modulated in response to polymicrobial sepsis. However, there were no differences in frequencies of *i*NKT or γδ T cells between MR1^-/-^ and WT mice (Supplementary Figure 3).

### Consistent with murine sepsis studies, MAIT cell frequency, activation, and effectors are altered in septic patients

We examined MAIT cells isolated from the blood of 33 septic patients (16 female and 17 male, mean age 59 +/- 2.7years, mean SOFA score 4.6) within 48 (± 24) hours of ICU admission for sepsis (day 1, D1). In 12 of these same patients (6 female and 6 male, mean age 60.5 +/- 4 years), we repeated MAIT phenotyping again approximately 90 days after their ICU admission for sepsis. We compared them to 21 age- and sex-matched healthy donors (HD) (10 female and 11 male, mean age 51 +/- 3.8 years). The identified causes of sepsis included pneumonia, urinary tract, intra-abdominal, and soft-tissue infections. Upon ICU admission, septic patients showed lower frequencies of MR1-tetramer^+^ MAIT cells within the T cell (CD3^+^) compartment compared to healthy donors (HD) (Figure 5, A and B). At 90 days after sepsis, MAIT cell frequencies were higher than upon ICU admission, with levels comparable to HD (Figure 5C).

**Figure 5.**
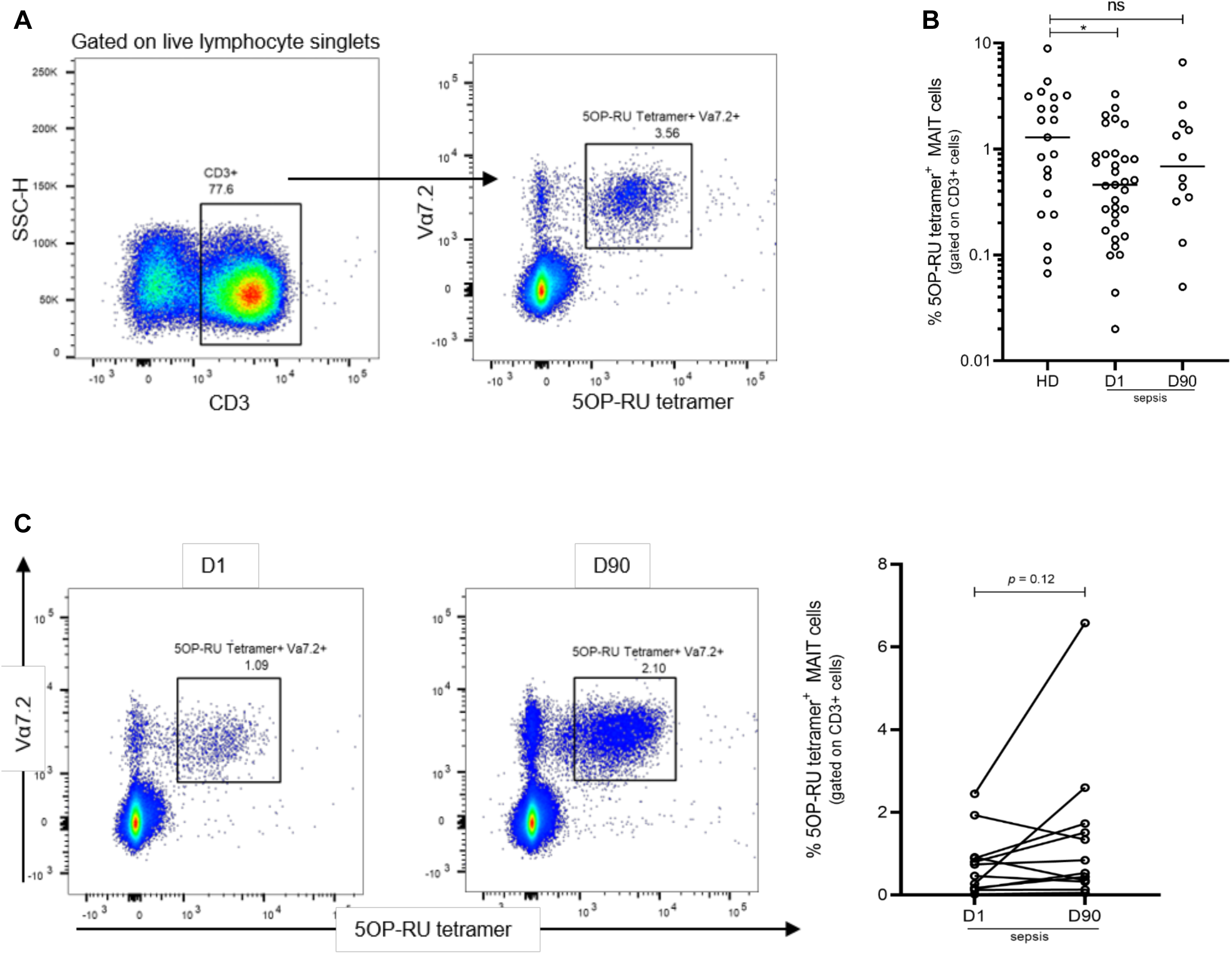
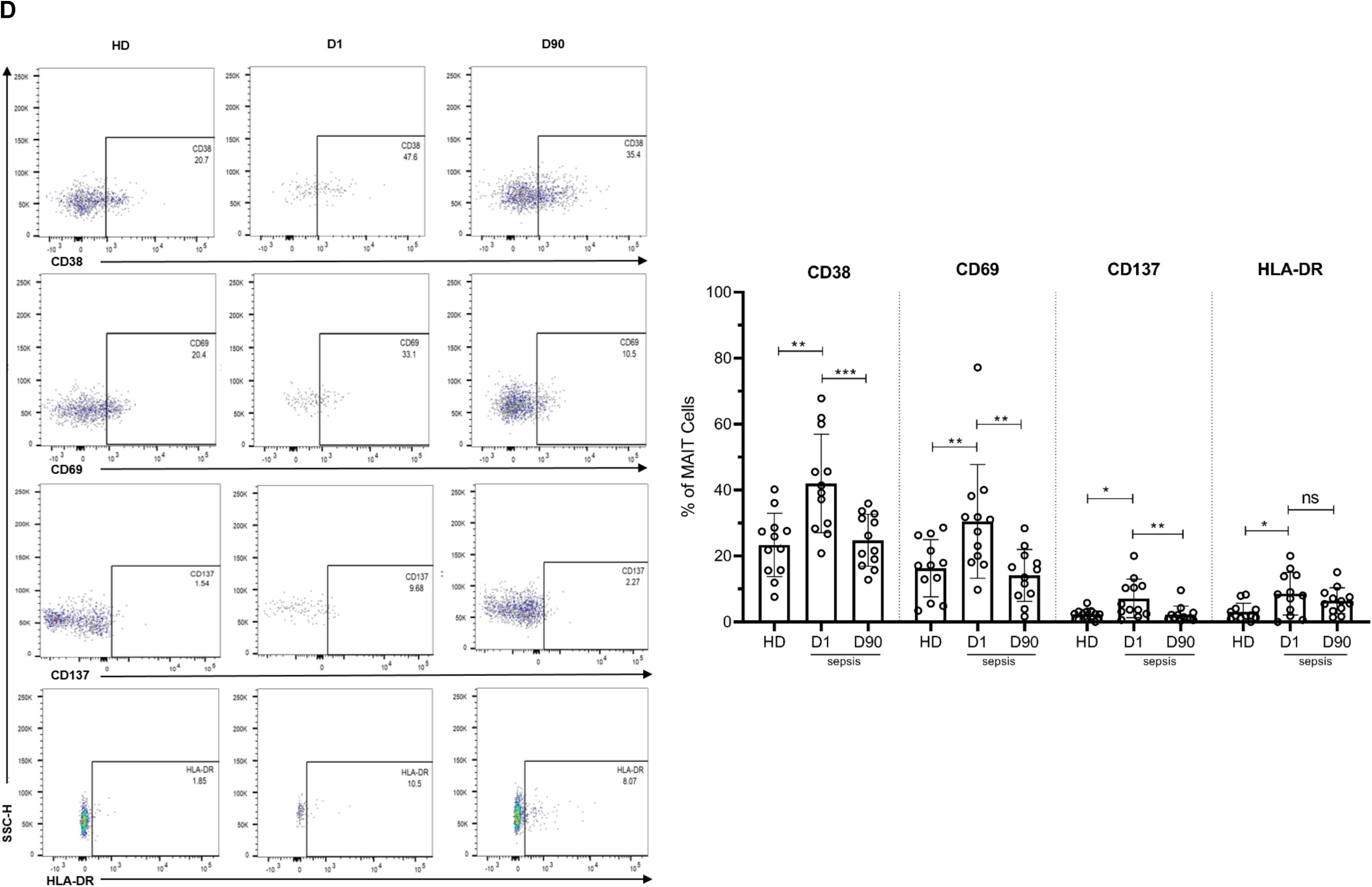
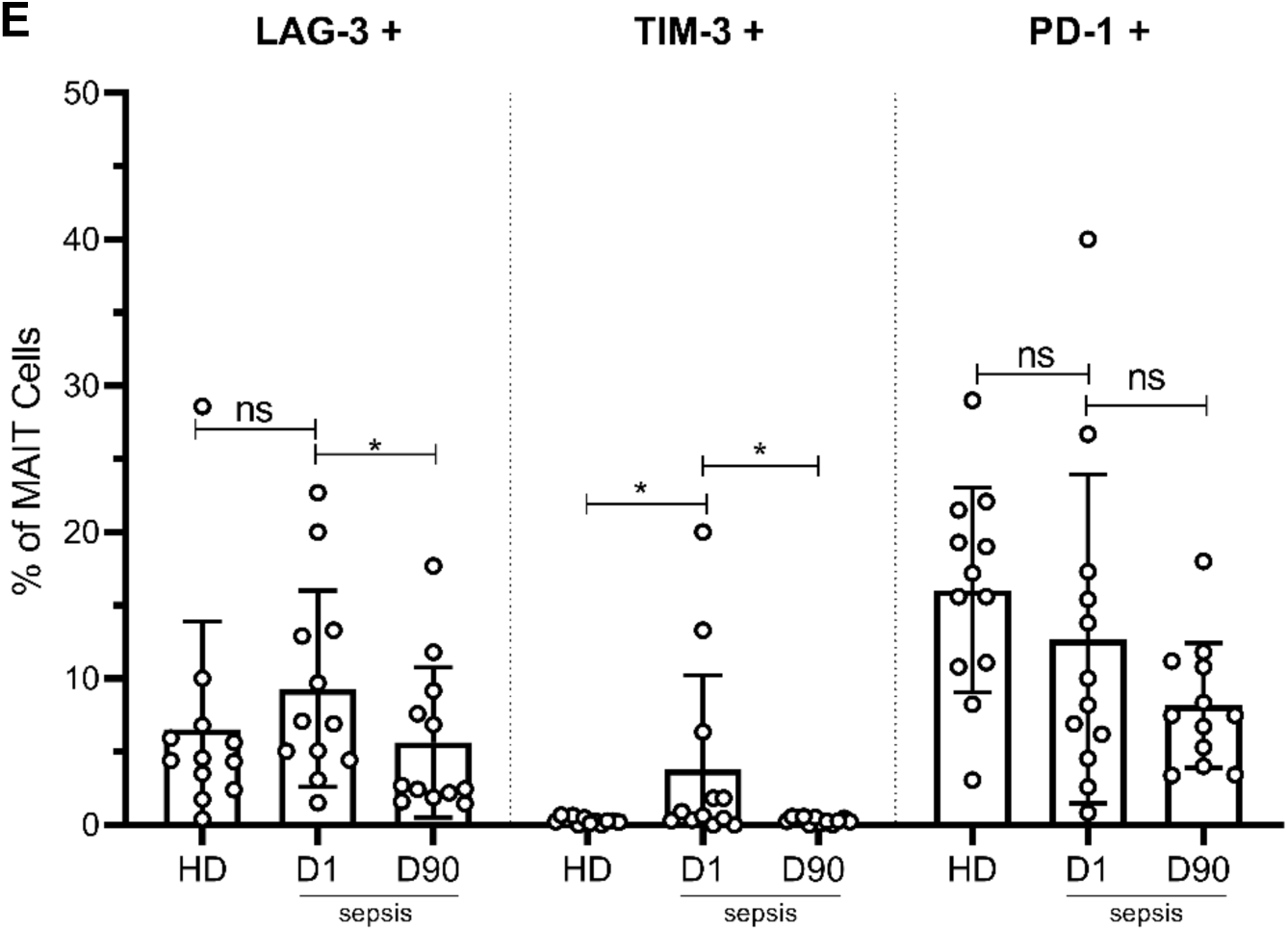
Altered frequency and phenotype of human MAIT cells in sepsis compared to healthy donors (HD). (**A**) Gating strategy for isolation of MAIT cells from PBMCs. (**B**) Percentage of MR1-tetramer^+^ MAIT cells in CD3^+^ T cells, (**C**) Representative flow plots and percentage of MAIT cells in paired Day 1 and D90 septic patients (n = 12 per group), (**D**) Representative flow plots and surface expression of activation markers as percentage of MAIT cells, (**E**) Surface expression of inhibitory receptors, as percentage of MAIT cells. Data were expressed as mean ± SD of two independent experiments. Mann-Whitney or Wilcoxon test was used. ***P < 0.001, **P < 0.01, *P < 0.05. NS, P > 0.05

Next, we evaluated markers of MAIT cell activation, exhaustion, and intracellular cytokine expression in a subset of surviving septic patients and for which paired samples at both Day 1 and Day 90 were available (n=12). Upon ICU admission (Day 1), the expression of MAIT cell activation markers, including CD69, CD38, and CD137, was significantly higher in septic patients compared to HD (Figure 5D). However, when assessed again in these same septic patients on Day 90, the expression of MAIT cell activation markers had returned to levels comparable to HD (Figure 5D). Upon ICU admission, we also found a higher expression of TIM-3 (P = 0.03), and LAG-3 (trend with P = 0.14), inhibitory receptors associated with T cell exhaustion, in MAIT cells of septic patients compared to HD (Figure 5E). As with MAIT cell activation markers, exhaustion markers LAG-3 and TIM-3 in septic patients at Day 90 had returned to levels similar to that of HD (Figure 5E). Of note, PD-1 expression was lower (but not statistically significant, P = 0.11) in septic patients compared to HD (Figure 5E).

We then examined MAIT cell effector functions by analyzing the intracellular expression of MAIT-associated inflammatory cytokines (IFN-γ, IL-17 and TNF-α) and granzyme B after *ex vivo* TCR stimulation using *E. coli*. Upon ICU admission, MAIT cells from septic patients showed significantly less IFN-γ production after *E. coli* stimulation, compared with HD. As with MAIT cell activation and exhaustion markers, IFN-γ production returned to levels comparable with HD at Day 90 (Figure 6A). No significant differences were observed in IL-17 (Figure 6B), TNF-α (Figure 6C) and granzyme B (Figure 6D) expression in MAIT cells among the groups.

**Figure 6.**
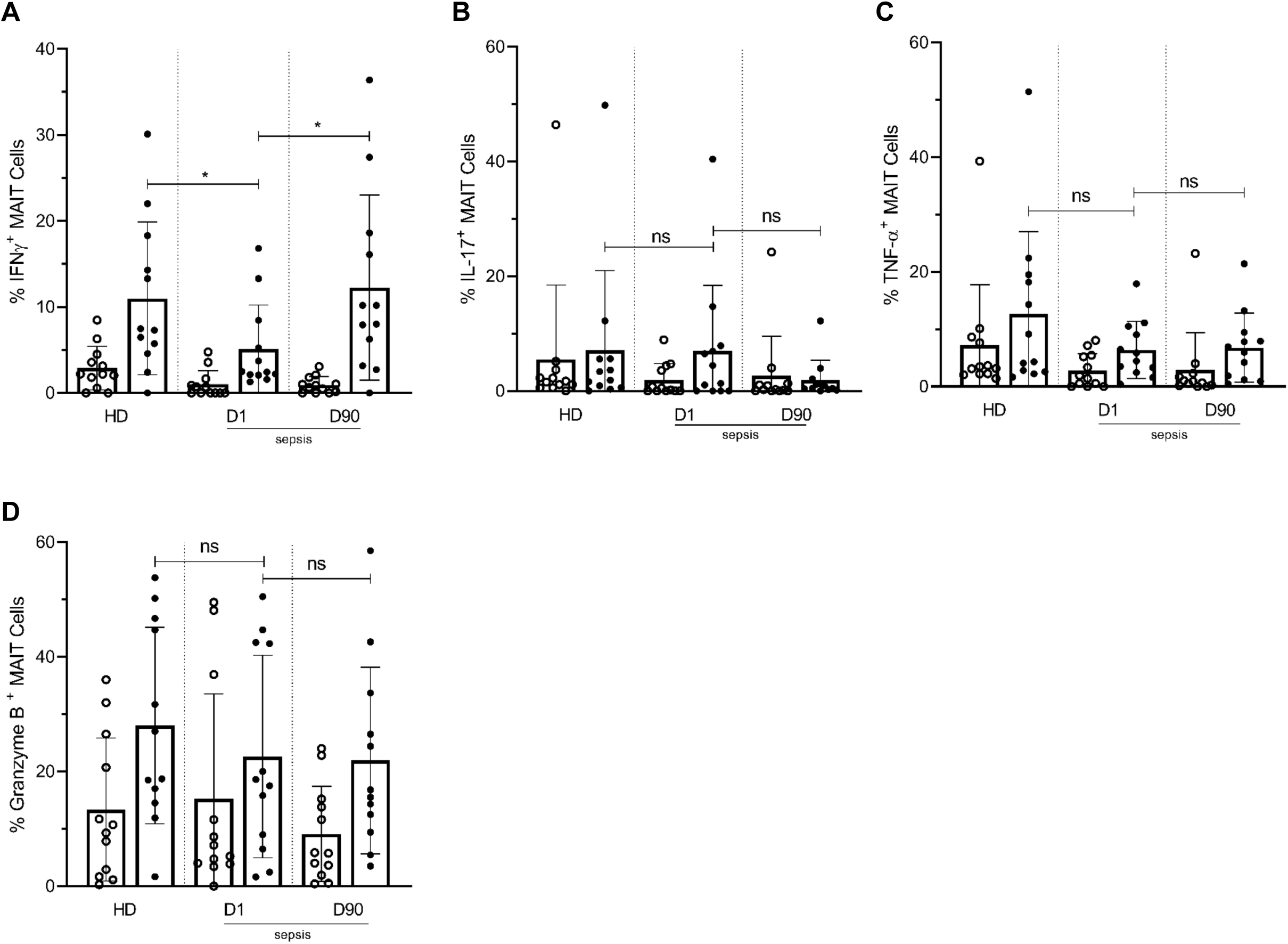
Lower frequencies of IFNγ^+^ MAIT cells in day 1 septic patients compared to HD and day 90 paired septic patients. PBMCs obtained from septic patients and healthy donors (n = 12 per group) were stimulated with *E.coli* at moi of 10 and intracellular expression of (**A**) IFN-γ, (**B**) IL-17, (**C**) TNF-α and (**D**) granzyme B by MAIT cells were analyzed using flow cytometry (open circle shows unstimulated and closed circle shows stimulated data). Data were expressed as mean ± SD of two independent experiments. Mann-Whitney or Wilcoxon test was used. ***P < 0.001, **P < 0.01, *P < 0.05. NS, P > 0.05.

Overall, these data demonstrate that during clinical sepsis, MAIT cell frequencies are lower, highly activated, and less capable of mounting IFN-γ responses upon stimulation. Moreover, at 90 days following sepsis, MAIT cell frequencies, activation, and IFN-γ responses have returned to levels comparable with matched HD.

## Discussion

Recent reports provide evidence that MAIT cells play an important role in antibacterial responses (29, 30). Despite this, very little is known about this unconventional T cell in sepsis. Our study reveals major alterations in activation, function, and dysfunction of MAIT cells in both clinical and experimental models of sepsis. We show that during clinical sepsis, MAIT cells are highly activated, lower in frequency, and have an altered functional phenotype evidenced by reduced IFN-γ cytokine expression. Moreover, at 90 days after sepsis, these changes in MAIT cells have resolved, as compared to matched healthy subjects. Using relevant animal models of sepsis, we show similarly that tissue-resident MAIT cells are dysfunctional during sepsis. Moreover, our experimental data indicate that MAIT cells are protective during sepsis, as a deficiency of MAIT cells resulted in significantly higher bacterial burden and mortality. These results indicate not only that MAIT cells are dysfunctional during sepsis, but also that MAIT cells may contribute to protective host responses and therefore outcomes during sepsis.

In both a polymicrobial model of sepsis (*i.e.* CLP) and a monomicrobial model using a clinical isolate of ExPEC, we found that MAIT-deficient (MR1^-/-^) mice have higher mortality, and that this is associated with a higher bacterial burden. We also found that these mice had lower levels of tissue cytokines (IFN-γ, TNF-α, IL-17, GM-CSF) than that in WT mice following induction of sepsis. We hypothesize that MAIT cells plays a protective role against sepsis pathology by regulating cytokine expression and maintaining a balance in immune response following sepsis to minimize tissue injury. As MAIT cells are capable of bridging the innate and adaptive immune systems, it has also been postulated that their actions are mediated in part by their interaction with macrophages or monocyte-derived dendritic cells. Interestingly, we found that in sepsis, MAIT-deficient mice produce less GM-CSF and have lower frequencies of lung interstitial macrophages and monocytic dendritic cells compared to WT mice. These observations are concordant with previous work demonstrating MAIT cell promotion of early pulmonary GM-CSF production, and differentiation of inflammatory monocytes into moDCs during pulmonary infection (24). Further work to examine whether tissue-resident MAIT cells have protective effects against sepsis pathology is warranted.

Our finding of decreased MAIT cell frequencies during severe sepsis are consistent with a previous study demonstrating that MAIT cells are decreased in patients with severe sepsis, a decrease that was not seen in other T cell populations examined, including *i*NKT and γδ T cells, suggesting cell-specific, rather than global, depletion (16). To extend this work, we found that at approximately 90 days after sepsis MAIT cell frequencies had returned to similar levels observed in matched healthy donors. Furthermore, we found that MAIT cells express elevated levels of activation markers in septic patients, and that these markers also return to similar levels as in healthy donors following recovery from sepsis. The depletion of MAIT cells from circulation in septic patients is similar to what has been observed in other infections (31) (32) (33), and may be a result of apoptotic cell death, TCR internalization, or migration to peripheral tissues, although the underlying mechanisms remain to be elucidated.

Previous work had shown an association of MAIT cell frequency with protection against secondary infections in the ICU (16). In this study, we found that in Day 1 sepsis patients, *ex-vivo* stimulated MAIT cells had a depressed IFN-γ response compared to healthy donors and patients reassessed 90 days following admission for sepsis. A similar decrease in IFN-γ response was also seen in MAIT cells in the lung tissue of mice undergoing CLP compared to mice undergoing sham surgery. Together with an increase in expression of co-inhibitory receptors on human MAIT cells, our data suggest that during clinical sepsis, MAIT cells are dysfunctional. This may prevent MAIT cells from mounting an optimal response against microbial pathogens, a hypothesis consistent with our experimental data indicating that loss of MAIT cells results in increased bacterial burden and mortality during sepsis. Previous work has shown that superantigens from bacterial pathogens such as *Staphylococcus spp.* and *Streptococcus spp.* can directly trigger rapid activation of MAIT cells to mount IFN-γ, TNF-α, and IL-2 responses in an MR1-independent manner (32), and that MAIT cells primed by superantigens are exhausted and anergic to cognate antigens such as *E.coli*.

In conclusion, we demonstrate that MAIT cells undergo functional and phenotypic changes during clinical and experimental sepsis. We found that MAIT deficient mice have higher microbial burden, lower tissue cytokine production, and higher mortality during sepsis. Our data provide new insights into the potential protective role of MAIT cells, and their contribution towards tissue-specific cytokine responses, during sepsis. Defining MAIT cells as a critical therapeutic target during sepsis, and defining the mechanisms that regulate MAIT cell function, will potentially enable the development of novel translational strategies targeted to enhance MAIT cell functionality to improve outcomes in sepsis.

## Methods

### Mice

10-14 weeks old male C57BL/6J wild type mice were obtained from Jackson laboratories and MR1^-/-^ mice, on a BL/6 background, were obtained from Siobhan Cowley (US FDA). All mice were bred in pathogen-free facility at the University of Utah. The animals were kept at a constant temperature (25°C) with unlimited access to pellet diet and water in a room with a 12 h light/dark cycle. All animals were monitored daily and infected animals were scored for the signs of clinical illness severity as previously described (20). Animals were ethically euthanized using CO_2._

### Mouse models of sepsis

C57BL/6 mice were anesthetized with ketamine/xylazine (100 mg/kg and 10 mg/kg, i.p., respectively), and sepsis was induced by cecum ligation and puncture (CLP) as previously described (Hubbard et al., 2005). After surgery, the animals received subcutaneous sterile isotonic saline (1 mL) for fluid resuscitation. Sham-operated mice were subjected to identical procedures except that CLP was not done. The animals were followed for 5 days after the surgical procedure for determination of survival. Bacterial loads were determined by serial dilution and plating of lung homogenates and blood on LB agar plates.

To examine monomicrobial sepsis, mice were injected intraperitoneally with 200 μl of PBS containing ∼1×10^6^ cfu of Extraintestinal pathogenic *Escherichia coli* (ExPEC) clinical strain F11 (34). Bacteria (F11) was grown from frozen stocks in static modified M9 media at 37°C for 24 hours prior to use in the sepsis model. Mice were monitored for 5 days for determination of survival.

### Lung mononuclear cell isolation

For lung digestion and preparation of single cell suspensions, lungs were perfused using sterile phosphate-buffered saline. After removal, the lung dissociation protocol (Miltenyi Biotec, Bergisch Gladbach, Germany) was performed using the mouse Lung Dissociation Kit (Miltenyi Biotec) and the gentleMACS Dissociator (Miltenyi Biotec) as per manufacturer’s instructions. After dissociation, cells were passed through a 70 µm cell strainer and washed with RPMI with 10 % FBS. Red blood cells were lysed with red blood cell lysis buffer. Lung mononuclear cells were then washed twice in RPMI with 10 % FBS before use in subsequent experiments.

### Tetramer and surface-staining of lung single cell suspensions

From each group of animals, 1-2 million cells aliquots were prepared and to exclude dead cells from analysis cells were first stained with the fixable viability dye eFluor™ 780 (eBioscience) for 15 min. at room temperature (RT). Cells were incubated with anti-mouse CD16/CD32 Fc Block antibody (BD Biosciences, USA), for 20 min at 4 °C. Cells were then stained for 30 min at RT with appropriately diluted PE conjugated 5-OP-RU-loaded species-specific MR1-tetramers or α-GalCer (PBS-44)–loaded CD1d tetramer conjugated to APC, anti-CD3-FITC (Biolegend), anti-CD161-BV510 (Biolegend), anti-CD49b-BV711 (BD), anti-TCRγδ-PE-Cy7 (Biolegend), anti-TCRβ-BV421 (Biolegend), anti-CD45R-PE-Cy5 and anti-CD44-BV650 (Biolegend). To evaluate different antigen presenting cells, after Fc Block incubation, cells were also surface stained with anti-CD45 AF700 (Biolegend), anti-CD11b-FITC (Biolegend), anti-CD11c-PE Cy-7 (BD), anti-Siglec-F-BV711 (BD), anti-CD64-BV605 (Biolegend), anti-CD24-PE (Biolegend), anti-Ly-6G-PerCP-Cy5.5 (Biolegend), anti-CD103-BV510 (Biolegend), anti-CD86-APC (Biolegend), anti-Ly6C (PE dazzle 594) and anti-MHC II BV421 (Biolegend) for 30 min at 4 °C. Total 10^6^ gated events per sample were collected using Fortessa flow cytometer (Becton Dickinson, San Diego, CA), and results were analyzed using FlowJo 10.4.2 software.

### Quantitation of MAIT cytokine transcripts by real-time PCR

MAIT cells (CD3^+^ CD44^+^ TCRβ^+^ MR1-5-OP-RU tetramer^+^) and non-MAIT (MR1-5-OP-RU tetramer^-^ TCRβ^+^) CD3^+^ T cell populations were sorted into 350 µl RLT plus buffer from the Qiagen Allprep DNA/RNA micro kit. Nucleic acid extraction was performed according to manufacturer’s instructions (Qiagen Allprep RNA/DNA). cDNA was synthesized using SuperScript VILO cDNA Synthesis Kit and qRT-PCR analysis on *Ifng, Il17a, Gzmb* and *Prf1* genes was performed with specific TaqMan primers and TaqMan Fast Advanced Master Mix (Applied Biosystems). The mRNA levels of specific genes were determined by the relative standard curve method, normalized against housekeeping ribosomal protein L32 levels. ΔCT was calculated as CT_gene_ – CT_housekeeping_ and ΔΔCT was calculated as ΔCT_CLP_ – ΔCT_sham._

### Mouse cytokines

Following lung dissociation, cells were passed through a 70-μm nylon mesh. Cells and debris were removed from the suspension via centrifugation at 300 x g for 10 min and supernatant were collected and stored at −80^°^C for later use. Lung cytokine levels were assessed from the supernatant samples via LEGENDplex (mouse inflammation panel 13-plex; BioLegend) kit per manufacturer’s instructions. Serum samples were 2-fold diluted and cytokine levels were assessed using same kit as per manufacturer’s instructions. Cytokine levels were acquired using a FACSCanto II flow cytometer (BD Biosciences, San Jose, CA), and analyses were performed using LEGENDplex data analysis software (BioLegend).

### Human Subjects

Patients admitted to an academic medical intensive care unit (ICU) with a primary diagnosis of severe sepsis or septic shock were prospectively enrolled within 48 (±24) hours of ICU admission. Sepsis was defined using the consensus criteria (35, 36) at the time this study was actively recruiting, defined as suspected or confirmed systemic infection and organ dysfunction as defined by a sequential organ failure assessment (SOFA) score ≥2 above baseline. Septic shock was defined as sepsis and an elevated lactate > 2mmol/L or the need for vasopressors. Blood samples were collected from septic patients upon study enrollment (Day 1, after ICU admission for sepsis) and again in the same subjects ∼90 days after enrollment (Day 90). Venous blood was collected and centrifuged over a Ficoll-Hypaque density gradient using the standard protocol to isolate peripheral blood mononuclear cells (PBMCs), which were either cryopreserved or used directly. Samples from healthy donors were collected and processed following the same procedure.

### Analysis of MAIT cytokine expression by flow cytometry

For phenotypic analysis of MAIT cells, PBMCs were thawed from −80°C and were stained for surface markers: anti-CD137-Alexa Fluor 700 (Biolegend), anti-CD3-BUV395 (BD biosciences), anti-CD8-PE-Cy5.5 (Molecular Probes), anti-CD4-BUV496 (BD Biosciences), anti-Vα7.2-BV711 (Biolegend), anti-LAG3-BV785 (Biolegend), anti-CD25-BV650 (Biolegend), anti-PD-1-BV605 (Biolegend), anti-CD161-BV510 (Biolegend), anti-CD69-PE-Cy5 (Biolegend), anti-HLA-DR-FITC (Biolegend), anti-TIM3-BV421 (Biolegend), anti-CD38-PE-Cy7 (Biolegend), and anti-human MR1 5-OP-RU Tetramer (NIH Tetramer Core Facility). PBMCs that were cultured and stimulated with 1100-2 *E. coli* (*E. coli* Genetic Stock Center (CGSC), Yale University) at multiplicity of infection (MOI) of 10 for 18 hours were stained for surface markers: anti-human MR1 5-OP-RU Tetramer (NIH Tetramer Core Facility), anti-CD3-BUV395 (BD biosciences), anti-CD8-PE-Cy5.5 (Molecular Probes), anti-CD4-BUV496 (BD Biosciences), anti-Vα7.2-BV711 (Biolegend), anti-LAG3-BV785 (Biolegend), anti-CD25-BV650 (Biolegend), anti-PD-1-BV605 (Biolegend), anti-CD161-BV510 (Biolegend), anti-CD69-PE-Cy5 (Biolegend) and intracellularly stained for: anti-IL-17A-FITC (Biolegend), anti-IFN-γ-PE-Cy7 (Biolegend), anti-TNFα-ef450 (Molecular Probes), anti-Granzyme B-Alexa Fluor 700 (Biolegend). All samples were acquired using a 5-laser BD LSRFortessa II flow cytometer (BD Biosciences) and analyzed using FlowJo software v10 (Tree Star, Inc. Ashland, Oreg).

### Statistics

Comparisons between independent groups were performed with unpaired t test. The Mann-Whitney *U* test was used for comparison of continuous variables between two groups and results of lung cytokine analysis were presented as mean ± standard deviation. For comparisons of more than two groups, one-way ANOVA with Tukey’s multiple comparison test was used.

For comparisons of healthy subjects with day 0 or day 90 septic patients Mann-Whitney *U* test was used and for comparisons of paired day 0 and day 90 samples Wilcoxon test was used. GraphPad Prism 8.3.0 software was used for all statistical analysis and *p* < 0.05 was considered statistically significant.

### Study approval

Each patient or a legally authorized representative provided written, informed consent. The institutional review board approved this study. All animals were maintained and experiments were performed in accordance with The University of Utah and Institutional Animal Care and Use Committee (IACUC) approved guidelines (protocol # 18-10012).

## Author contributions

S.T., C.P.A., R.A.C., J.S.H., M.T.R, and D.T.L. conceived and designed experiments; T.B., E.M., enrolled human subjects; S.T., D.L., C.P.A., C.V.A., E.M., A.T. and R.A.C. performed experiments and analyzed data; S.T. and D.T.L. wrote the initial draft of the manuscript; and D.L., C.P.A., M.A.M., R.A.C., J.S.H., and M.T.R., provided manuscript feedback and advice.

## Acknowledgements

This research was supported by the National Institutes of Health (AI130378 to D.T.L., HL092161, AG040631, and AG048022 to M.T.R., TL1TR002540 to D.L., T32 HG008962 to C.P.A.], and a University of Utah 3i Initiative Seed Grant (to D.T.L, M.T.R., J.S.H.). We would like to thank all the study subjects who participated in the study. We would like to thank Michael C Graves and Alexandra Heitkamp for help in laboratory. We would also like to thank the staff of the University of Utah Flow Cytometry Core and the Office of Comparative Medicine.

---

**Supplementary Figure 1.**
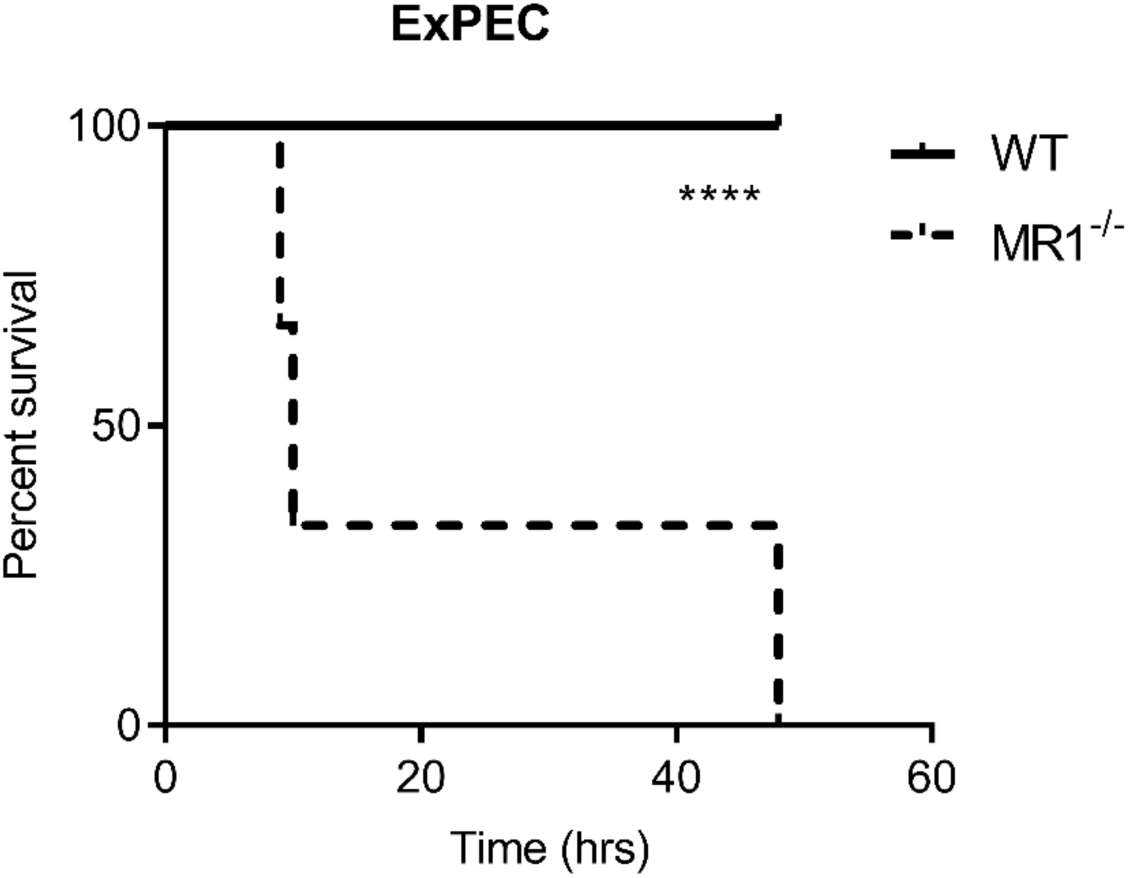
MR1^-/-^ mice had significantly higher mortality from ExPEC sepsis, compared to WT mice. Graph shows survival curves after intraperitoneal injection with 1×10^6^ cfu of Extraintestinal pathogenic *Escherichia coli* (ExPEC) F11 strain.

**Supplementary Figure 2.**
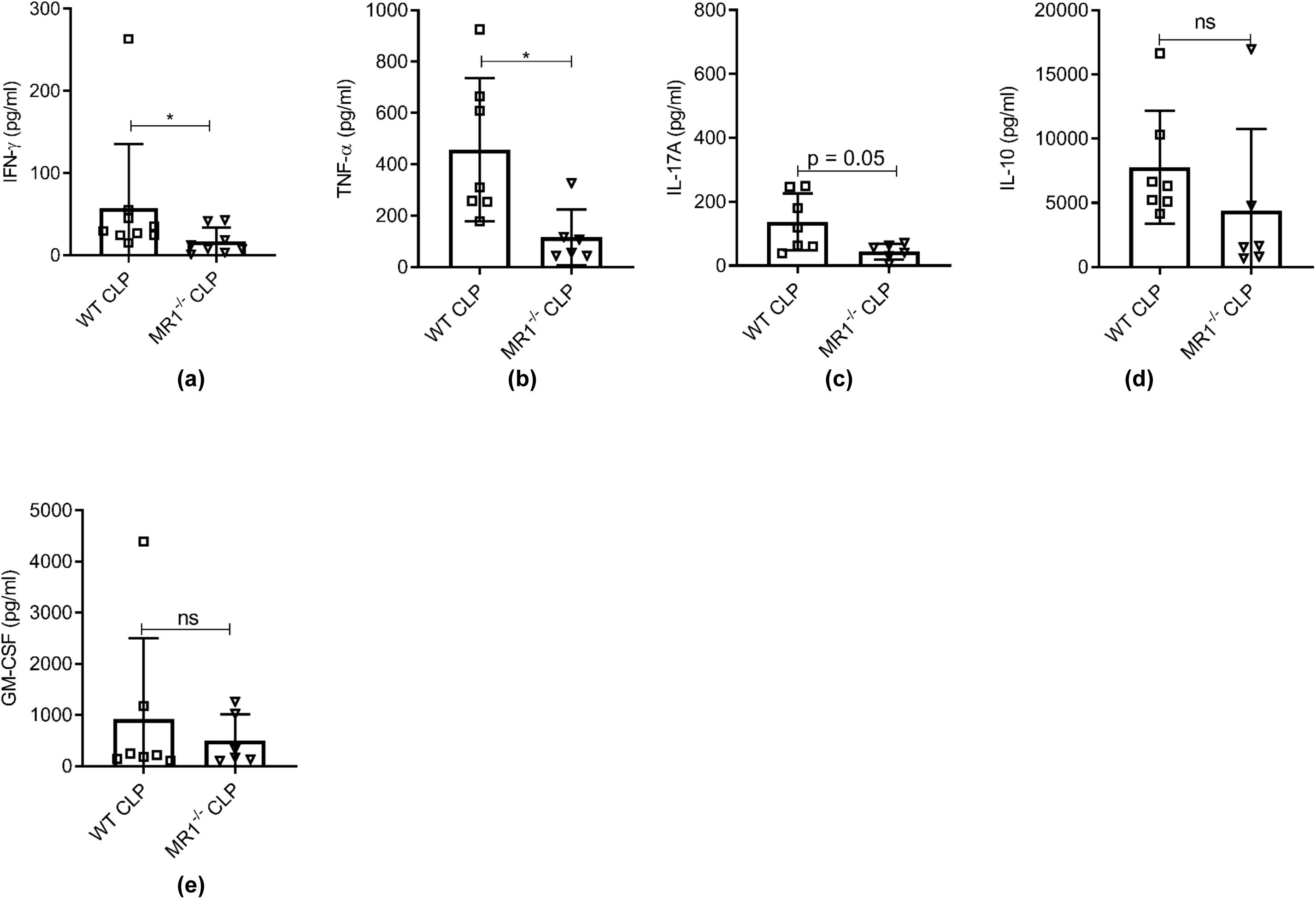
MR1^-/-^ mice had reduced IFNγ and TNFα but similar levels of other cytokines in serum at 18 h post CLP. Levels of (a) IFN-γ, (b) TNF-α, (c) IL-17A, (d) IL-10, and (e) GM-CSF were assessed in serum of WT and MR1^-/-^ mice at 18 h post CLP using a bead-based immunoassay kit. Bar represent mean cytokine level ± SD. The graphs represents average of two independent experiment (WT n = 7-8, MR1^-/-^ n = 6-7). The statistical significance was determined by the nonparametric Mann–Whitney test.

**Supplementary Figure 3.**
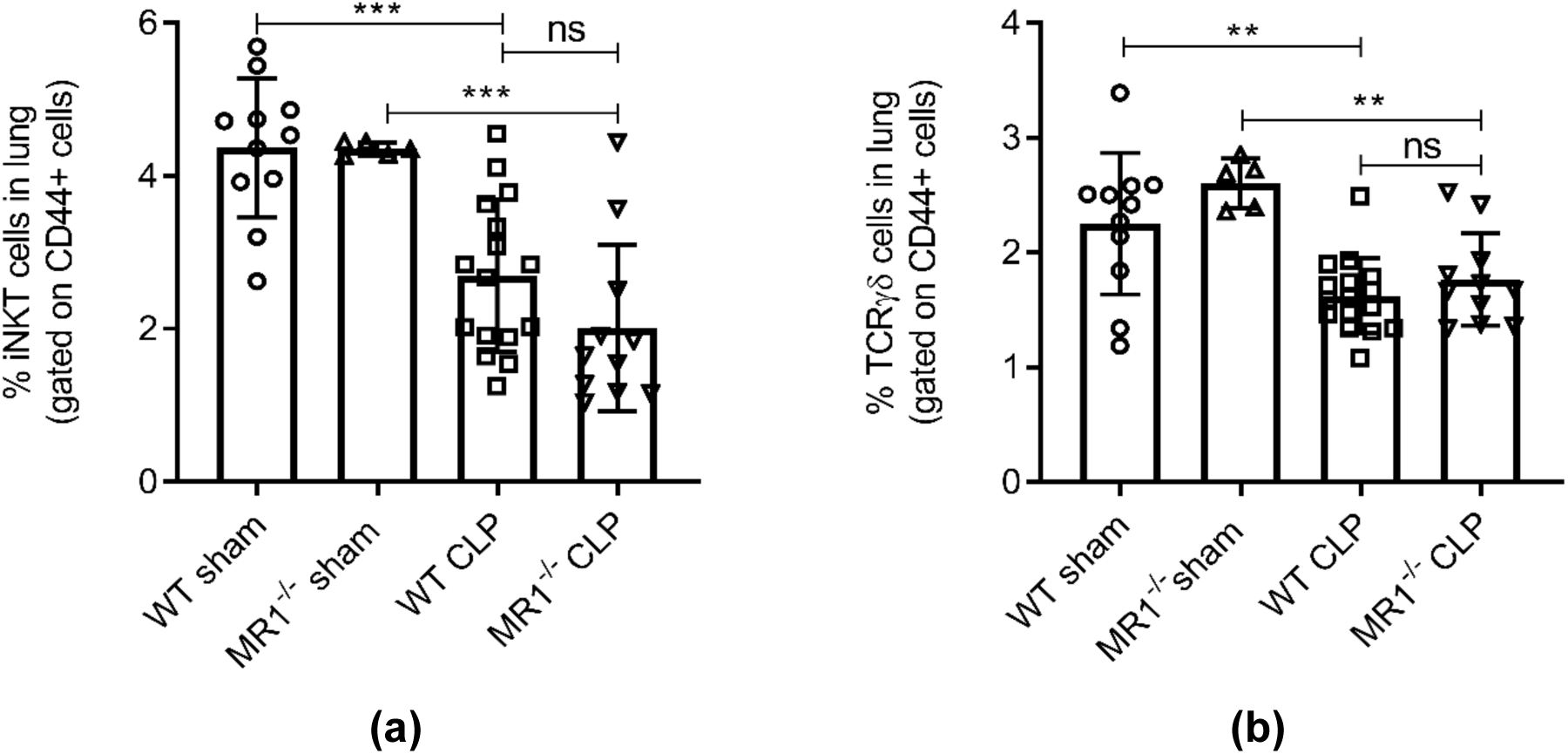
Frequencies of other unconventional T cells such as invariant natural killer T cells (iNKT) and γδ T cells post CLP. Lung cells were obtained from WT and and MR1^-/-^ mice at 18 h post CLP or sham operation, stained for surface markers and analyzed using flow cytometry. Percentage frequencies of (a) *i*NKT cells and (b) γδ T cells were compared between the groups using one-way ANOVA with Tukey’s multiple comparison test.

